# A dual-mode mobile phone microscope using the onboard camera flash and ambient light

**DOI:** 10.1101/162008

**Authors:** A. Orth, E. R. Wilson, J. G. Thompson, B. C. Gibson

## Abstract

Mobile phone microscopes are a natural platform for point-of-care imaging, but current solutions require an externally powered illumination source, thereby adding bulk and cost. We present a mobile phone microscope that uses the internal flash or sunlight as the illumination source, thereby reducing complexity whilst maintaining functionality and performance. The microscope is capable of both brightfield and darkfield imaging modes, enabling microscopic visualization of samples ranging from plant to mammalian cells. We describe the microscope design principles, assembly process, and demonstrate its imaging capabilities through the visualization of unlabelled cell nuclei to observing the motility of cattle sperm.

## Introduction

The rapid advancements in imaging capabilities of consumer mobile phones over the last decade have made such devices attractive for point-of-care and resource-poor microscopy applications. Microscopy-enabled mobile phones have been used for a variety of purposes including malaria diagnosis(1), sperm tracking(2–4) and water quality assessment(5). These mobile phone microscopes come in a variety of form factors with a range of working principles. An early design was comprised of a standard microscope objective interfaced to the mobile phone camera via a lens tube and eyepiece(6); a later iteration more resembles a miniaturized benchtop microscope, with a 3D printed stage and chassis(7). A simpler, lower resolution mobile phone microscope consists of an additional camera lens along with the integrated camera lens to form a unity magnification imaging system(8). Together with an external light emitting diode (LED) and stage, this forms a transmission mode brightfield microscope, with enough magnification to image red blood cells. This basic optical design consisting of a magnifying lens and an external LED also forms the basis of the ultra-low-cost Foldscope, made primarily from origami paper(9).

Another class of mobile phone microscopes are lens-free devices that image via holography(2,5,10). With these devices, the sample is placed directly onto the image sensor. The sample is then illuminated by an external light source in a particular geometry so as to create a series of holograms, which are captured by the image sensor. Subsequent image processing translates the raw holograms into images. Advantages of this approach are increased resolution and light collection efficiency since there is no lens to limit the numerical aperture. Lens-free techniques are also amenable to tracking in 3D since the holograms carry 3D information. However, the image is not viewable in real-time, and often requires processing on a powerful desktop computer or in the cloud. More importantly, the user must disassemble the camera module itself in order to remove the lens and place the sample directly on the image sensor. Cleaning the sensor after use is also not practical. These challenges are a hurdle for wide scale adoption of lens-free mobile phone techniques.

Despite the assertion that mobile phone microscopes are simple, low-cost tools for use outside the lab, most mobile phone microscopes require extra components, most notably external illumination modules. Two published exceptions are a lens-free device that uses the sun as an illumination source(5), and brief report that describes the use of diffuse reflection from a slide holder placed behind a sample(11). Aside from these two publications, every mobile phone microscope described in the literature features an externally powered LED light source. Battery-powered LEDs present an issue if and when they run out of energy as finding a replacement battery or recharging the battery may not be possible in remote settings. External LEDs also add extra bulk and assembly complexity to a system that is meant to be as compact and simple as possible. Ideally, a mobile phone microscope would take advantage of the integrated flash found in nearly every modern mobile phone, obviating the need for external lighting and power. The difficulty in using the built-in camera phone flash is that the flash is offset from the camera by typically a few mm, and is pointing in the same direction as the camera.

Reflection mode microscopy is not possible in this configuration since the flash does not illuminate a sample located near the camera’s entrance aperture; transmission illumination mode is also not possible as it requires the light source and camera to be on opposite sides of the sample along the optical axis. Using the integrated flash of the microscope appears to require additional mirrors and lenses to turn and condense the illumination light back onto the sample. This in turn would necessitate additional optical components (adding cost and bulk), and consequently would defeat the purpose of using the internal flash.

In this work we describe a 3D printed microscope add-on clip that enables transmission brightfield and darkfield microscopy on a mobile phone without any externally powered light source or additional illumination optics. For brightfield transmission mode, our design takes advantage of the integrated phone flash together with diffuse reflection in a similar manner to that previously noted(11). Unlike in this previous report, our 3D printed device itself has the necessary geometry to create diffuse transmission illumination without employing an external diffusely reflective object behind the sample. Moreover, darkfield imaging is made possible by designing the clip so that ambient light only can illuminate the sample via internal reflection within the sample glass slide. As a result, we can observe samples that are nearly invisible under brightfield operation due to low absorption or refractive index contrast, such as cells in media.

Because our design requires no external power or light sources, it is particularly robust, making it ideal for use in remote areas and field work. Our microscope requires only a single assembly step (inserting the lens into the 3D printed clip), avoiding more complicated assembly involving electrical hardware and multiple 3D printed parts. The simplicity not only makes it easy to set up and use but also helps to drive down cost when assembly costs are taken into account.

In this paper, we outline the design and operational principles of our mobile phone microscope followed by optical characterization and application examples. We have also made the Solidworks and STL files for 3D printing freely available to enable users around the world to print their own microscope clip.

## Materials and Methods

### Microscope clip design

Our mobile phone microscope design consists of a 1x magnification imaging system that is created by placing a mobile phone camera lens (exterior to the mobile phone) in front of the mobile phone’s existing internal mobile phone camera module(8). This configuration resembles a classic infinite-conjugate microscope design that is the basis of modern optical microscopes(12). The exterior mobile phone camera lens is friction-fit into a recess of the clip (“lens” in Fig. 1a) and plays the role of objective lens, while the camera lens inside the phone takes on the role of the tube lens. If the sample is placed one focal length in front of the objective lens, an image is formed one focal length behind the tube lens. Mobile phones are designed to image near infinity by default, meaning that the standard position of image sensor is typically one focal length behind the tube lens. In such an infinite conjugate system, the optical magnification is the ratio of the focal length of the objective to tube lens - 1x in our case. However, because the pixel size of mobile phone cameras is so small (1.22μm for iPhone 6s), 1x magnification nevertheless results in microscopic resolution. As noted previously, the advantage of using a mobile phone camera lens as the objective is that these lenses are very well corrected for aberrations, thus yielding images far superior to those captured with a more simple optic such as a ball lens(8). Moreover, due to mass manufacturing, mobile phone camera lenses are inexpensive, especially given that they have multiple lens elements. One can extract the camera lens from an iPhone camera module purchased online for $ 15 AUD(13). This cost can be further reduced by purchasing camera lenses themselves in bulk direct from the manufacturer.

**Figure 1.**
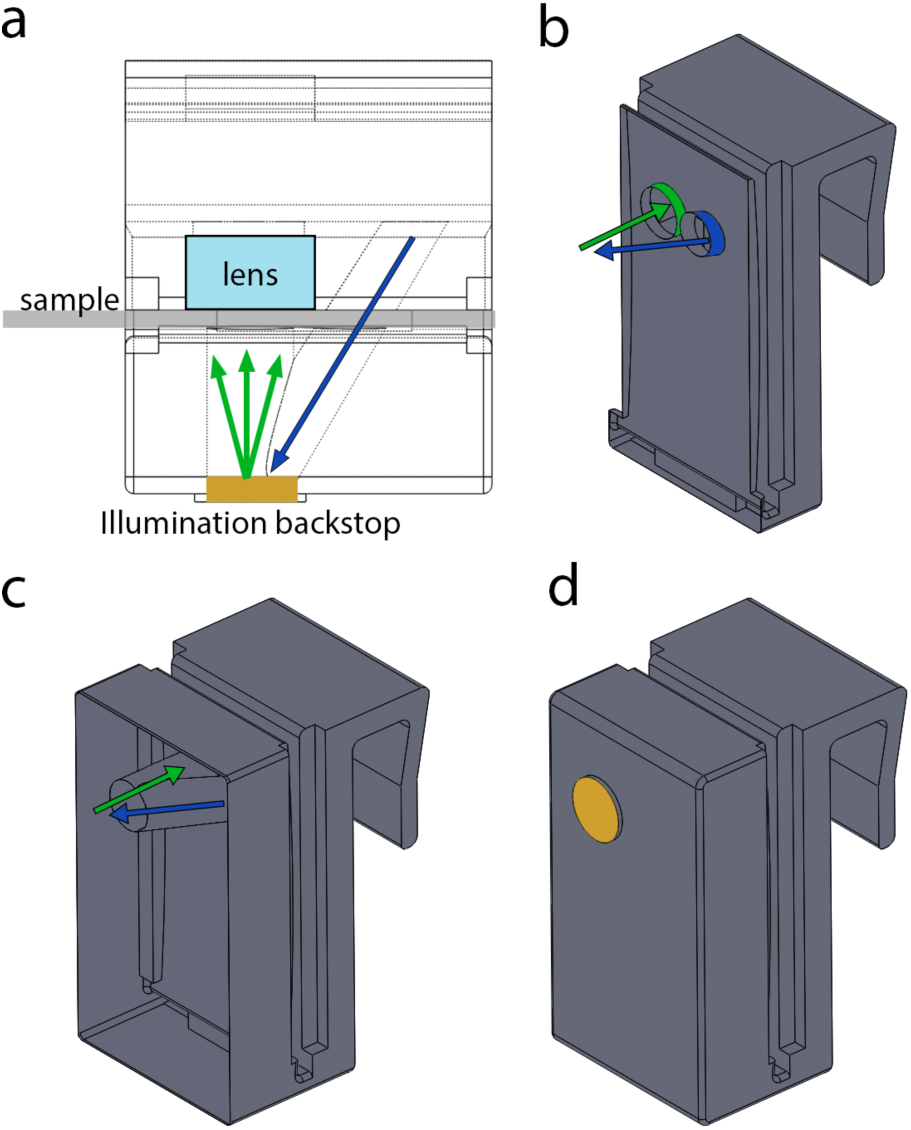
Renderings of the microscope clip Solidworks CAD file. a) A wireframe overhead view of the clip, showing the illumination tunnels. The blue arrow denotes light exiting the flash, and green arrows indicate diffusely reflected light from the resin backstop (gold). The sample slide is shown in grey, and the lens label indicates the location of the external objective lens when placed inside the objective lens recess. b) A cross-section view of the clip showing the illumination tunnels, the outside of which are highlighted in blue and green for pre- and post-diffuse reflection tunnels, respectively. Light exiting the flash is first confined to the pre-diffuse reflection tunnel (blue). Light then travels back through the post-diffuse reflection tunnel (green) after which it illuminates the sample. The direction of light in each tunnel is indicated by the coloured arrows. c) A cross - section of the microscope clip showing the entirety of the illumination tunnels (no longer highlighted in colour). The tunnels intersect at the back of the clip at which point light from the flash encounters a diffusely reflective backstop of cured resin. As in (c), the coloured arrows indicated the direction of light propagation in each tunnel. d) A rendering of the entire microscope clip. The diffusely reflective backstop is highlighted in gold.

The novelty of our device lies in the illumination design. Instead of employing an external LED, we use the internal mobile phone flash. In order to use the flash in transmission geometry, we design a microscope clip with internal illumination tunnels, as shown in Fig. 1. The entrance of the tunnel is placed at over the camera flash. Light from the camera flash travels through the first tunnel (blue in Fig. 1a,d), reflects diffusely off of the end of the tunnel (gold in Fig. 1c) and then travels back into another tunnel that is aligned to the optical axis of the objective lens and camera module (green in Fig. 1a,d). After being diffusely reflected, light from the camera flash illuminates the sample in transmission (green arrows in Fig. 1d). A similar approach has been used to trans-illuminate samples in endoscopy, where one does not have access to the rear of the sample(14). Despite losses from reflections within and at the end of the illumination tunnel, there is still ample light at the sample plane for imaging. This is perhaps unsurprising considering that the camera LED is designed to illuminate objects at distances much larger (>1m) than the path length in our microscope (~1cm). Another convenient feature of this approach is that the illumination results from a diffuse reflection. Ideally, brightfield transmission microscopy is performed with Kohler illumination(15), where light travelling in all possible angles admitted by the numerical aperture (NA) of the condenser hits every point in the sample. Despite the lack of a condenser lens, our geometry approaches this condition because diffuse reflections create light travelling in all directions. In our microscope clip, the effective illumination NA (0.23) is defined by the area over which the diffuse reflection occurs (a circle of radius 2.63mm) and the distance between the diffuse reflector and the sample (11mm). This illumination NA matches the NA of the imaging objective (f/2.2), just as in a standard brightfield microscope.

When operating the mobile phone microscope, the user can choose to turn the flash on or off. Turning the flash on results in a transmission brightfield image as described. However, with adequate ambient lighting, an image is still created when the flash is off, despite the lack of direct sample illumination. This image is the result of light trapped via (total) internal reflection in the microscope slide being scattered into the objective lens by the sample. In this configuration, light cannot land on the image sensor unless it is scattered by the sample, resulting in a dark background(16). Darkfield imaging modalities such as this one are particularly useful for observing samples that do not absorb strongly or that are nearly index matched to their surroundings.

### Fabrication and assembly

We print our microscope clip using a Formlabs Form 1 3D printer. The microscope clip design file is converted from STL to the native Formlabs format and then uploaded to the printer via the standard PreForm software. For acceptable contrast in darkfield mode, it is necessary for the microscope clip itself to be opaque. As a result we print the microscope clip in black resin (Formlabs GPBK02). Printing in white or grey resin severely degrades darkfield imaging performance. For optimal print speed, we print without supports and select a layer thickness of 0.1mm. Printed clips are then rinsed in IPA as per the suggested Formlabs protocol(17). After rinsing, the microscope clip is let to dry and is then put under sunlight for 1-2 hours to post-cure (post-cure optional).

After drying and post-curing, the objective lens must be inserted into the microscope clip. The objective lens fits into a rectangular recess designed to hold the lens in place via friction. The objective lens is inserted into the clip from the front as shown in Figs. 2a-d. Squeezing the microscope slide holder sets the objective lens into place at approximately the right position. If needed, the objective lens position can be adjusted by pushing it further into the recess using tweezers. Note that the objective lens should be oriented in the opposite direction to the internal camera (tube) lens. That is, the surface of the objective lens that was previously facing the sensor should be facing the sample.

**Figure 2.**
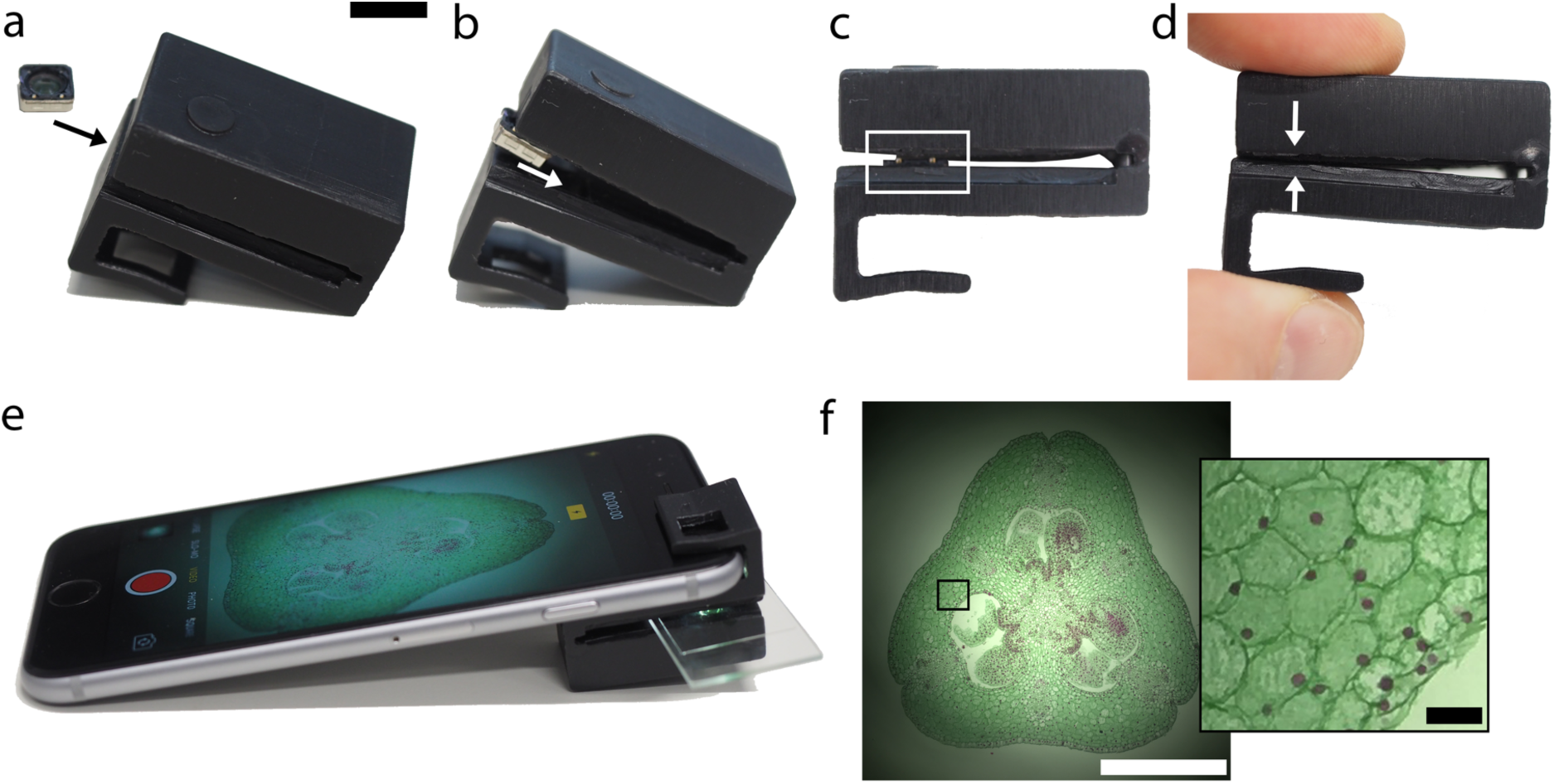
Mobile phone microscope assembly process. a) Insert mobile phone camera lens (objective lens) into microscope clip as shown. Make sure that the side of the lens assemble that originally faced the image sensor now faces the sample (faces away from the camera). Scale bar approx. 1mm for (a)-(d). b) Push objective lens further into the clip until it fits into the recess. c) Gently push the objective lens assembly into the recess. This can be done with tweezers or by hand. The white boxed region shows the objective lens assembly sitting in the friction-fit recess. d) Gently squeeze microscope clip so that the opposite sides of the slide holder come into contact. This pushes the objective lens assembly into its final position in the microscope clip recess. e) Insert sample slide and attach the clip to an iPhone 6s as shown. The objective lens fits directly over the iPhone back camera. Open the iPhone camera app (or other 3^r^d party camera app), switch to video mode and activate the flash to view the sample in brightfield mode. In this example, the sample is Lilium ovary (Southern Biological). f) Brightfield image of Lilium ovary using “Photo” mode with flash. Scale bar is 1mm. Inset: Magnified image of boxed region. Scale bar is 50μm.

Once the objective lens is in place, the clip is fitted over the mobile phone such that the objective lens is directly over the internal camera module (Fig. 2e). Samples mounted on microscope slides can be inserted into the clip as shown in Fig. 2e. The native iPhone camera app enables either brightfield (with flash on) or dark field imaging (flash off). An example of a brightfield image of a Lilium ovary (Southern Biological) acquired with the microscope is shown in Fig. 2f.

## Results and Discussion

### Resolution characterization

The optical magnification of the mobile phone microscope is equal to the ratio between the focal length of the lenses. We use an iPhone 5s back camera lens paired with the internal iPhone 6s back camera lens, which both have an f/2.2 and focal length 4.15mm, giving 1x optical magnification. The pixels on the iPhone 6s camera are on a 1.22μm pitch, suggesting a Nyquist-limited resolution of 2.44μm. However, the colour Bayer filter together with the camera’s internal demosaicing reduces the achievable resolution because only one colour is sampled per pixel. Though the optical magnification is fixed at 1x, the digital zoom and imaging mode of the iPhone affects the apparent pixel size. At native 1x digital zoom in “Photo” mode of the native iPhone camera app, we measure an effective pixel size of 1.22μm, as expected. At full digital zoom in “Photo” mode, the effective pixel size is reduced five-fold to 0.24μm. In “Video” mode, pixel sizes were measured to be 2.19μm and 0.73μm at 1x and 3x digital zoom, respectively. Note that at 1x digital zoom in “Video” mode, the iPhone undersamples the image sensor, making it necessary to use digital zoom in order to maintain full spatial resolution.

We measure the microscope resolution by imaging USAF-1951 resolution targets. We employ two types of targets, one consisting of 2μm thick photoresist (Nanoscribe IP-Dip) features on a quartz slide (phase target, Fig. 3a & c), the other with transparent features on a chrome-coated glass slide (chrome target, Fig. 3b). Transparent objects, such as the photoresist features on the phase target are sometimes harder to resolve than opaque features, as they do not produce as much intensity contrast. Despite this expectation, we find that the resolution of both the chrome and phase targets are essentially equal, at 4.48μm and 4.38μm, respectively. In darkfield, the phase target is easily visible under ambient room light. The resolution, however, is slightly worse than in brightfield mode, with the smallest resolvable grating having a pitch 5.60μm.

**Figure 3.**
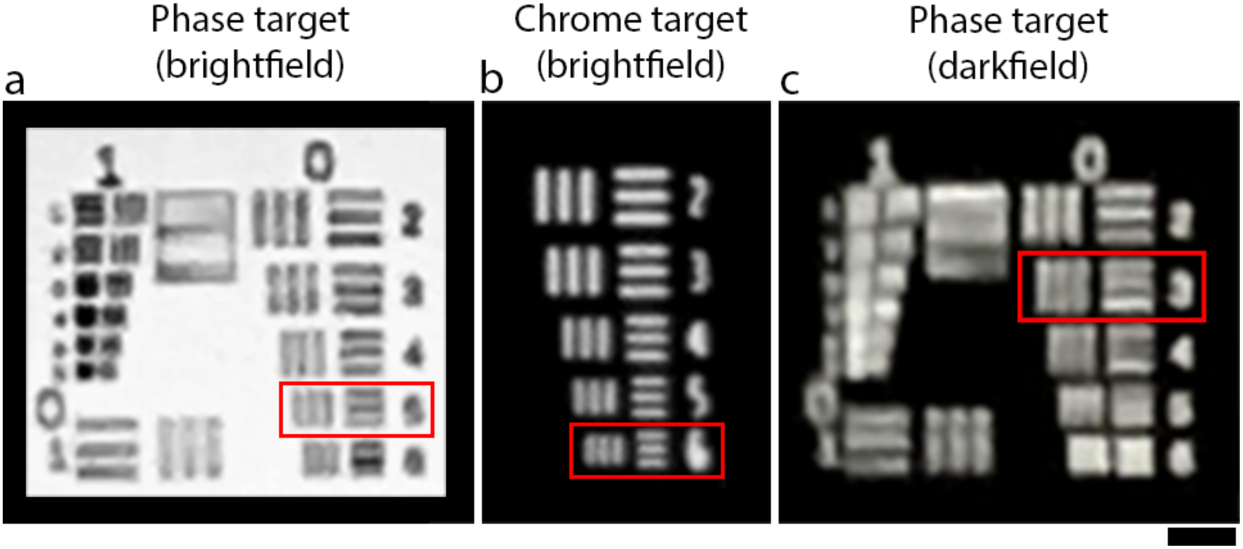
Images of resolution targets taken by the mobile phone microscope under various contrast mechanisms. Red boxes indicate the smallest resolved gratings in each case. a) A phase object target where features consist of 2μm thick bars of photoresist (n=1.48) on glass in air (n=1). The smallest resolved grating has pitch = 4.48μm. The light source is the phone flash. b) A portion of group 7 gratings on a chrome mask test target (a binary opaque/transparent mask). The light source is the phone flash. The smallest resolved grating has pitch = 4.38μm. c) The same phase target as in (a), but imaged in darkfield mode. The phone flash is turned off. Ambient roomlight is the illumination source. The smallest resolved grating has pitch = 5.60μm. Scale bar 20μm. All images recorded in “Photo” mode at 1x digital zoom.

Improved optical resolution could be achieved by using a shorter focal length objective lens. For example, the front camera lenses on smartphones typically have focal lengths in the range of 2-3mm, which could improve resolution up to two-fold over the 1x magnification system presented here. Even shorter length ball lenses or microlenses(18) are of potential interest, though these elements pose problems in terms of significant field aberrations(4,9).

### Cell Culture

The advantage of using darkfield illumination becomes apparent when imaging transparent objects such as cells in a nearly index-matching medium such as water. In brightfield transmission, the refractive index contrast within the cell and between the cell and its surroundings produces almost no intensity contrast. In darkfield, however, the illumination light is trapped inside the glass slide, mounting medium and coverslip, and can only escape if scattered. This leads to a dark background with bright features, which is the ideal situation for observing minimally absorbing, nearly index matched objects. The brightfield transmission image of a Caco-2 cell culture is shown in Figs. 4a & c. The cells are hardly visible due to low contrast, and it is unclear how many cells are actually present. Towards the edge of the FOV, where the effective illumination NA is lower, cells become visible, but barely so before the signal drops off significantly due to vignetting. A striking improvement in contrast is seen in a darkfield image of the same FOV (Fig. 4b).

**Figure 4.**
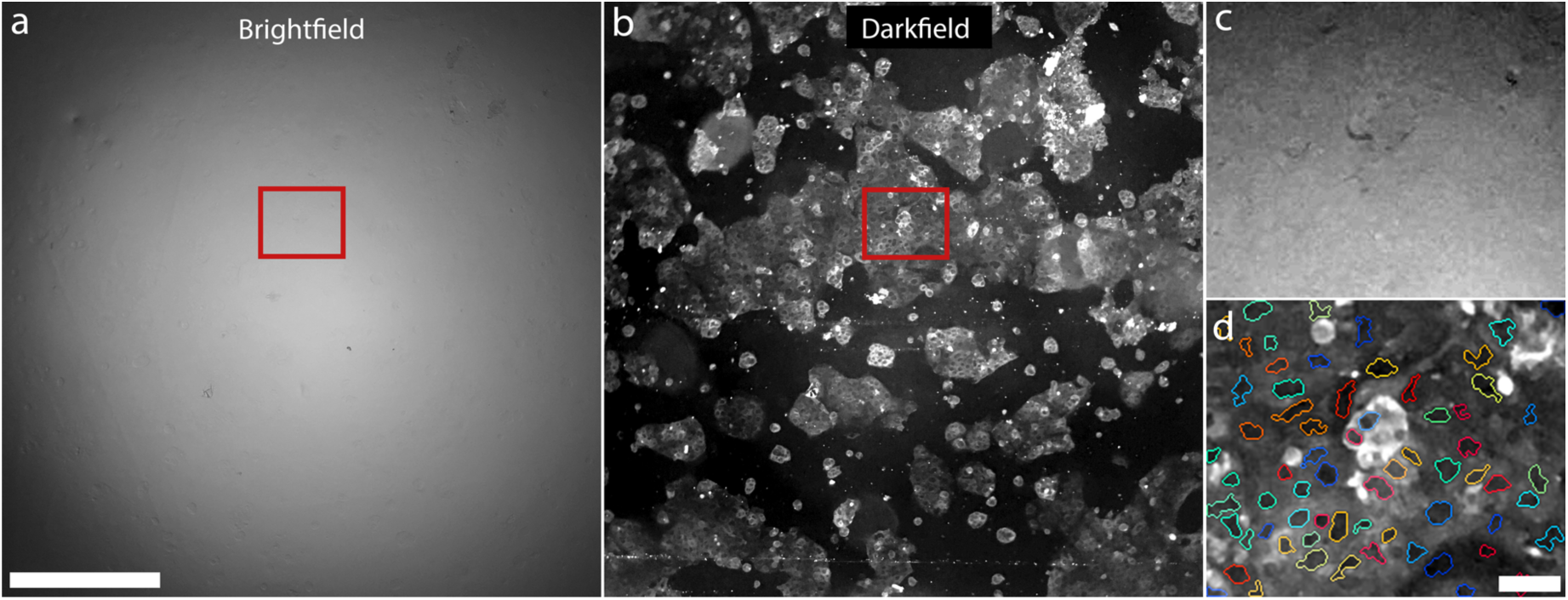
a) Brightfield image of unstained CaCO2 cells. The cells are nearly index-matched in the mounting medium, generating little contrast in brightfield. The light source is the phone flash. Scale bar 500μm. b) The same sample as (a), but imaged in darkfield. The phone flash is off, and sunlight is the illumination source. Cells appear with high contrast against a dark background. Cells nuclei are clearly visible as dark centres within bright cytoplasm. c) A magnified view of the red box in (a). Contrast has been enhanced for improved visibility. d) Magnified view of the red box in (b). Cell nuclei appear as dark circular/oblong features within bright cytoplasm. Cell nuclei are identified using an automated custom MATLAB nucleus finding algorithm, and outlined in a different colour for each cell nucleus. Only cells that scatter enough to saturate the detector are visible in the brightfield image in (c). Scale bar 50μm. All images recorded in “Photo” mode at 1x digital magnification.

Though the cells are nearly invisible under brightfield illumination, the darkfield image shows that the cell culture is in fact highly confluent. A careful comparison between Figs. 4c & d indicates that only the very strongly scattering cells that saturate the camera detector in darkfield produce enough contrast to be visible in brightfield. Not only are unlabelled cells visible in darkfield, but cell nuclei are clearly visible in a magnified view of a region of the FOV (Fig. 4d). The cell cytoplasm appears brighter than the nucleus likely because of all of the fine cellular features inside the cytoplasm acting as scattering centres. Using a custom MATLAB script(19), we show that the contrast between the nuclei, cytoplasm and the surrounding background is enough to enable basic cell counting, without resorting to fluorescent dyes or histopathology stains.

### Sperm Motility

Dynamic samples can be observed with the mobile phone microscope using the camera setting “Video” on the iPhone 6s. We test the feasibility of live cattle sperm quality assessment both in brightfield and darkfield modes on our mobile phone microscope. In brightfield, dark oval shaped spots corresponding to the sperm heads are visible. Figure 5a shows the first frame of a 21-second movie (Supplemental Movie 1) of cattle sperm swimming freely between a microscope slide and coverslip, recorded with our mobile phone microscope. In order to visualise the trajectories of all the sperm in the FOV, we construct an image where the sperm images for each frame in the movie are superimposed. The sperm images for each frame are then color-coded by hue so that one can follow the sperm trajectories in time (Fig. 5b, Supplemental Movie 2). From this image, one can identify differences in motility patterns (eg. Circular, forward progressive). This analysis gives a quick qualitative indication of the health of the semen sample, whereas a more quantitative picture can be obtained via tracking data.

**Figure 5.**
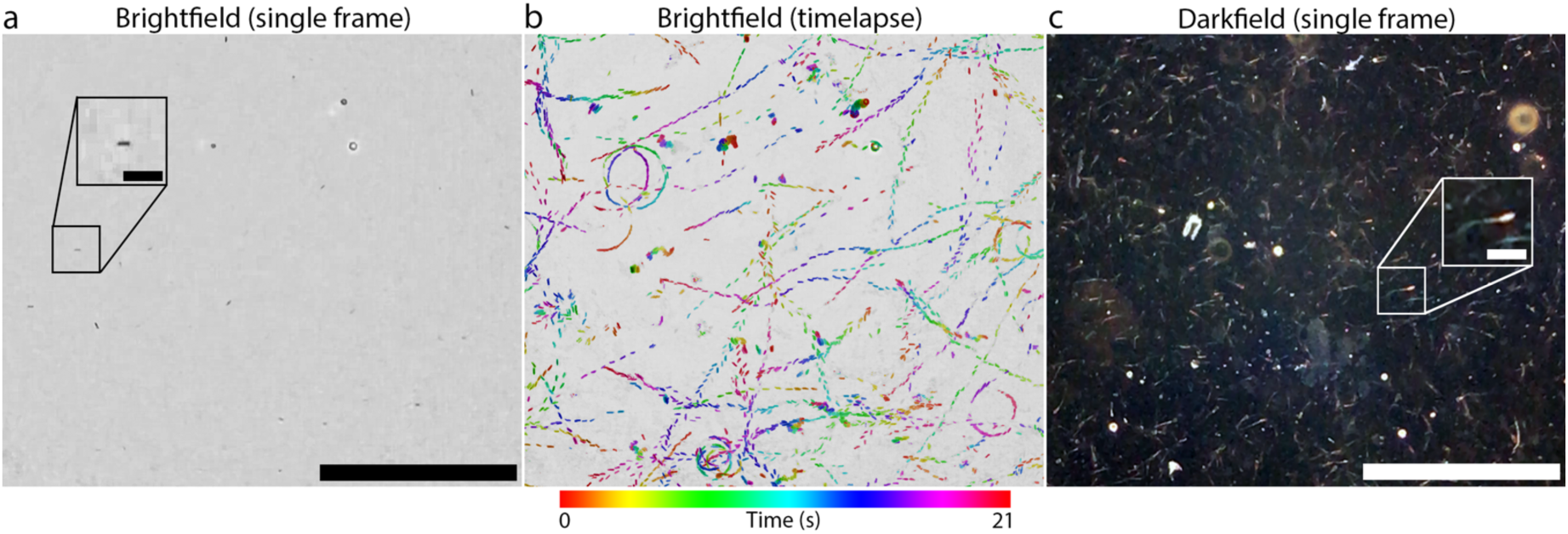
a) The first frame of motility tracks of sperm under brightfield illumination, recorded on our mobile phone microscope. Original movie is provided as Supplementary Movie 1. Scale bar is 300μm. Image is illumination corrected to compensate for vignetting. Inset: 2x magnified image of the boxed region containing a single sperm (head), which appears as a small elliptical dark spot. Inset scale bar 30μm. b) A representation of the entire brightfield movie from (a). Sperm locations are shown in colour with the hue changing through time. Colour bar shown below. Both circular and straight trajectories are visible. This figure is alternatively available as Supplementary Movie 2, where the colour-coded tracks appear over time. c) The first frame of a video of motile cattle sperm under darkfield illumination (sunlight). Scale bar 300μm. The field of view is different from (a) and (b). Inset: 2x magnification of small boxed region, showing a single cattle spermatozoan. The sperm head and tail are visible. Inset scale bar 30μm. All images recorded in “Video” mode at 3x digital magnification.

Sperm imaging can also be achieved with much higher contrast in darkfield mode with solar illumination. Figure 5c is the first frame of an 11-second movie (Supplemental Movie 3) of a cattle semen sample (frozen/thawed) mounted between a microscope slide and coverslip. In this imaging modality, the tails of the sperm are clearly visible in addition to the sperm heads. One drawback of the high contrast in darkfield imaging is that other scattering structures in the semen sample (lipid aggregates from semen extenders, other seminal debris, etc…) also contribute the image, potentially confounding sperm tracking algorithms. Ideal tracking performance may require further sperm purification steps as is standard in computer-aided sperm analysis(20).

## Conclusions

We have designed a simple mobile phone microscope that takes advantage of the integrated illumination available with nearly all smartphone cameras. Our design requires no additional illumination optics, reducing cost and assembly complexity. The microscope is useable after one assembly step and requires only one extra component: a readily available mobile phone camera lens. With this design we demonstrated both brightfield and darkfield microscopic imaging, including the visualization of cell nuclei in unlabelled cells and live cattle sperm. This device has the potential to be used as a general microscopy platform for a wide range of applications from biological fieldwork to microfluidic lab on a chip monitoring.

## Acknowledgements

This work has been supported by the ARC Centre of Excellence for Nanoscale BioPhotonics (CE140100003) and MicroNano Research Facility (MNRF) at RMIT University. B. C. G acknowledges the support of an ARC Future Fellowship (FT110100225). J. G. T. acknowledges the support of an NHMRC Research Fellowship (1077694).

